# Varietal Evaluation of Coffee Genotypes in Subtropical Nepal

**DOI:** 10.64898/2026.10.28.762398

**Authors:** Aditi Bhandari, Anup Bhatt, Sajan Gaudel, Suyasha Pudasaini, Asmita Khanal, Durga Devkota

## Abstract

Coffee (*Coffea arabica* L.) is an emerging high-value crop in Nepal, but national productivity remains far below genetic potential, partly due to the lack of locally validated variety recommendations. This study evaluated and compared the morphological and yield-attributing characteristics of 19 Arabica genotypes at the Horticulture Research Station, Malepatan, Pokhara (848 m above sea level) from January to June 2026 using a Randomized Complete Block Design with three replications. Yellow Caturra recorded the highest total fresh cherry yield (5.34 kg/plant), followed by Caturra Amarillo (3.56 kg/plant), Catuai Vermelo (3.19 kg/plant), and Catuai Amarillo (3.10 kg/plant). Catuai Amarillo and Caturra Amarillo produced the most bearing branches (76 and 72, respectively), while Caturra Amarillo and Yellow Caturra had the most cherries per node (7.73 and 7.47). Pacamara and Selection-10 achieved the maximum 100-cherry weight (167.67 g), while Indonesia had the largest cherry diameter (13.38 mm). Catuai Amarillo had the highest total soluble solids (25.97° Brix), while Tekisic had the lowest (16.80° Brix). Coffee leaf rust severity increased from a mean of 7.49% at 30 days after incidence (DAI) to 37.58% at 120 DAI. Selection-10 was the least affected genotype across the severity assessments and recorded the lowest area under the disease progress curve (AUDPC; 25.03 rating-days), whereas Caturra Amarillo had the highest AUDPC (347.95 rating-days). Overall, Yellow Caturra showed the strongest yield performance, while Selection-10 combined low disease development with a high 100-cherry weight. These results indicate that genotype selection should consider productivity, quality-related traits, and CLR response together.

## Introduction

Coffee (*Coffea arabica* L.) is a high-value crop with environmental and economic importance, now cultivated across more than 41 districts in the mid-hills of Nepal, including Gulmi, Palpa, Syangja, Arghakhachi, Kavre, Sindhupalchowk, Lalitpur, Kaski, Tanahu and Lamjung **[1]**. The region’s climate, topography, and soil conditions are well suited for Arabica cultivation **[2]**, typically at an altitude of 800-1600 masl **[3]**, with agroforestry and shade-grown systems being used widely to cope with and maintain optimal shade to regulate temperature extremes, moisture stress, and wind exposure **[4]**. Coffee farming has proven promising, especially in the midhills of Nepal due to the presence of favourable soil with an appropriate microclimate **[5]**. Although it contributes to national GDP, it remains small at roughly 0.01% **[6]**. It delivers a higher return per unit area than staple cereals and many cash crops **[7]**.

Despite this potential, coffee production in Nepal continues to underperform. Nepal’s national coffee productivity stands at 107.96 kg/ha **[8]**, far below the global averages and yields demonstrated in research trials, where optimal plots have yielded up to 7.04 kg per plant at altitudes of 900–1,100 masl **[9]**. The gap is due to multiple constraints; the major constraint remained the continued cultivation of inappropriate or poorly adapted genotypes **[10,11]**. The vegetative and yield data derived from other countries may not be similar for Nepal’s conditions due to the variation in agro-ecological conditions. Locally selected genotypes including Chhetradeep, Arghakhachi Local, Kaski Local, and Syangja Special have not been systematically characterized in the published literature, leaving farmers without objective data on their comparative merit.

Genotype-by-environment interaction is a well-established constraint on the transferability of yield performance data across locations. The World Coffee Research International Multilocational Variety Trial (IMLVT), which evaluated 29 genotypes at 23 sites across 15 countries, demonstrated that genotype-by-environment interactions substantially affect both yield and CLR resistance expression, underscoring the necessity of local field evaluation rather than extrapolation from global data **[12]**. Comparable GXE effects have been reported in Ethiopia, where site-genotype interaction accounted for a significant share of yield variance **[13,14,15]**, reflecting that genotypes’ yield or resistance ranking in one region carries limited weight in another. For Nepal, the stakes extend beyond current production: up to 72% of existing coffee-growing areas are projected to become climatically unsuitable by 2050 **[16]**. The remaining land must be planted with genotypes proven to perform locally and with premium genotypes like Bourbon and Pacamara, which already command price premiums of 200–500% above commodity Arabica in international specialty markets **[17]**. This raises the cost of getting the wrong selection of genotype for the local area.

Yield potential alone is not sufficient for recommending any genotype, since disease pressure remains a major constraint in coffee production regardless of genotype’s yield potential. Coffee leaf rust, caused by *Hemileia vastatrix* **[18]**, is one of the most destructive diseases of coffee worldwide **[19]**. Under severe infection, CLR can reduce coffee yield by up to 70% and can cause complete foliar loss, with berry loss reaching 70% **[20]**. In Nepal specifically, shifts in rainfall patterns linked to climate change are reported to intensify the CLR intensity across coffee-growing regions **[21]**. This study evaluated the 19 *Coffea arabica* genotypes and compared yield performance, morphological characteristics, and coffee leaf rust severity assessed via disease incidence, severity scoring and area under disease progress curve (AUDPC) of 19 genotype of coffee in Kaski, Nepal with aim to identify best genotype suited to the region based on yield, quality and disease resistance.

## Materials And Methods

### Experimental materials and treatments

The experiment was conducted at Horticulture Research Station (HRS) Pokhara, Nepal, with a latitude of 28° 13’ 034” N, a longitude of 83° 58’ 27.8” E, and 848 masl. The city had a humid subtropical climate with mild temperatures because of its elevation **[22]**. The atmospheric conditions (temperature and precipitation) throughout the study period were obtained from the NASA Prediction of Worldwide Energy Resources (NASA POWER) database and aggregated into weekly values for graphical presentation.

### Parameters recorded

#### Observation of morphological, size-related, and yield-related traits of coffee

Morphological traits of coffee were observed by counting the number of bearing branches, number of non-bearing branches, nodes per branch, and cherries per node. Yield parameters were calculated by counting the number of cherries per plant, fresh cherry yield, and hundred-cherry weight. Cherry length and diameter were size parameters, measured using a digital vernier caliper with an accuracy of 0.01 mm.

### Observation of Total Soluble Solids (TSS) content and dry parchment and wet parchment of different genotypes of coffee

#### TSS of pulp

The Total Soluble Solids (TSS) of cherry juice was analyzed using a digital refractometer with an accuracy of ±0.2%. A total of 500 g of coffee cherries were weighed, which were then pulped. Pulped coffee cherries were packed into a plastic bag, creating anaerobic conditions.

#### Dry parchment and Wet Parchment

After harvesting, ripe cherries were immediately pulped, fermented for about a week, and washed to obtain wet parchment, which was then weighed. Coffee cherries of 500grams for each variety were selected. Wet parchment was then air-dried, and the resulting dry parchment was weighed.

#### Determination of status of disease severity and incidence-related parameters in various genotypes

Coffee leaf rust severity was assessed on each genotype using a visual score scale ranging from 0 to 10, following **[23]**, where a score of 0 indicated no visible symptoms and a score of 10 indicated maximum disease severity. Rust scoring was carried out by observing the total number of leaves on each upper, middle, and lower branch. For incidence assessment, leaves were counted separately from the upper, middle, and lower branches of each plant. Scoring was done after the onset of disease infection and followed for four months. For calculation of disease incidence, a similar protocol was followed as done in disease severity, and a similar assessment window was followed.

#### Statistical analysis

Prior to the statistical analysis, the underlying assumptions of general linear model-linearity, normality of the residuals (Skewness and Kurtosis), corrected link function, and homogeneity of variance (heteroscedasticity) were assessed using the Global Validation of the Linear Model Assumption (GVLMA) test **[24]**. The GVLMA test was run on the raw data and indicated a violation of the model assumption. Data were therefore transformed where necessary using log(x+1), where x represents the observed value and 1 is added to accommodate 0 values. GVLMA was reapplied to the transformed data, confirming that all the model’s assumptions were satisfied.

Using Microsoft Excel 365, data entry, compilation, and organization were accomplished. RStudio 4.3.1 was used to conduct an analysis of variance (ANOVA) test with the package “Agricolae” in the R environment, and mean separation was done by using Duncan’s Multiple Range Test (DMRT) method.

## Results And Discussion

The morphological and yield characteristics of the 19 genotypes of coffee were evaluated in order to evaluate the performance of these diverse local and introduced germplasms.

### Results Morphological traits

#### Number of bearing branches

The number of bearing branches varied significantly among coffee genotypes, with a grand mean of 45.13. Catuai Amarillo recorded the highest number of bearing branches (76.00), followed closely by Caturra Amarillo (72.00) and Hawai Kona (70.17). The lowest number of bearing branches was recorded in Pacas (19.00) and Kaski local (14.50), which were significantly similar to each other. The number of bearing branches ranges from 14.50 to 76.00.

#### Number of non-bearing branches

Significant variation was observed in non-bearing branches, which had a grand mean of 13.72. Chhetradeep (45.67) and Caturra Vermelo (37.83) were found statistically similar and recorded the highest count for non-bearing branches. The lowest counts were recorded in Caturra Amarillo (3.83), Bourbon Vermelo (4.83), and Catuai Amarillo (5.17), which were significantly similar.

#### Cherry per node

Regarding the cherries per node, which had a grand mean of 5.81, the variety Caturra Amarillo (7.73) recorded the highest, followed by Yellow Caturra (7.47). The minimum number of cherries per node was recorded in the genotypes Bourbon Amarillo (4.93), Kaski Local (4.20), Mundo Novo (4.53), Pacamara (5.13), and Pacas (4.37), which are statistically at par.

In the case of nodes per branch, no statistical variation was found, indicating that all the genotypes are statistically at par.

**Table 1.** Description of coffee genotypes evaluated in the experiment.

| Genotype | Notation | Genotype | Notation |
| --- | --- | --- | --- |
| Arghakhachi Local | T1 | Kaski Local | T12 |
| Bourbon Amarillo | T2 | Mundo Novo | T13 |
| Bourbon Vermelo | T3 | Pacamara | T14 |
| Catimor | T4 | Pacas | T15 |
| Catuai Amarillo | T5 | Selection-10 | T16 |
| Catuai Vermelo | T6 | Syangja Special | T17 |
| Caturra Amarillo | T7 | Tekisic | T18 |
| Caturra Vermelo | T8 | Yellow Caturra | T19 |
| Chhetradeep | T9 |  |  |
| Hawai Kona | T10 |  |  |
| Indonesia | T11 |  |  |

**Table 2.** Average morphological traits of coffee (*Coffea arabica* L.) genotypes in Kaski, Nepal, 2026.

| Treatment | Bearing Branches | Non-Bearing Branches | Cherry per Node |
| --- | --- | --- | --- |
| **Arghakhachi Local** | 50.83 (3.88)<sup>a</sup> | 5.50(1.87)<sup>de</sup> | 5.33<sup>bc</sup> |
| **Bourbon Amarillo** | 31.69 (3.29)<sup>ab</sup> | 13.00 (2.62)<sup>bcd</sup> | 4.93<sup>c</sup> |
| **Bourbon Vermelo** | 49.50 (3.92)<sup>a</sup> | 4.83 (1.76)<sup>e</sup> | 5.57<sup>abc</sup> |
| **Catimor** | 35.67 (3.44)<sup>ab</sup> | 25.50 (3.26)<sup>ab</sup> | 6.13<sup>abc</sup> |
| **Catuai Amarillo** | 76.00 (4.30)<sup>a</sup> | 5.17(1.82)<sup>e</sup> | 6.53<sup>abc</sup> |
| **Catuai Vermelo** | 59.00 (4.05)<sup>a</sup> | 30.83 (3.28)<sup>ab</sup> | 6.47<sup>abc</sup> |
| **Caturra Amarillo** | 72.00 (4.23)<sup>a</sup> | 3.83(1.57)<sup>e</sup> | 7.73<sup>a</sup> |
| **Caturra Vermelo** | 35.33(3.54)<sup>ab</sup> | 37.83 (3.66)<sup>a</sup> | 6.33<sup>abc</sup> |
| **Chhetradeep** | 54.50 (3.5 3)<sup>ab</sup> | 45.67 (3.70)<sup>a</sup> | 5.53<sup>abc</sup> |
| **Hawai Kuna** | 70.17(4.26)<sup>a</sup> | 7.83(2.18)<sup>cde</sup> | 5.57<sup>abc</sup> |
| **Indonesia** | 34.00(3.47)<sup>ab</sup> | 15.00(2.68)<sup>bc</sup> | 6.13<sup>abc</sup> |
| **Kaski Local** | 14.50(2.68)<sup>b</sup> | 6.83(2.01)<sup>cde</sup> | 4.20<sup>c</sup> |
| **Mundo Novo** | 40.33(3.68)<sup>ab</sup> | 8.00(2.06)<sup>cde</sup> | 4.53<sup>c</sup> |
| **Pacamara** | 33.17(3.51)<sup>ab</sup> | 5.83(1.90)<sup>de</sup> | 5.13<sup>c</sup> |
| **Pacas** | 19.00(2.72)<sup>b</sup> | 12.83(2.60)<sup>bcd</sup> | 4.37<sup>c</sup> |
| **Selection 10** | 35.00(3.48)<sup>ab</sup> | 7.50(2.12)<sup>cde</sup> | 6.40<sup>abc</sup> |
| **Syangja Special** | 35.17(3.57)<sup>ab</sup> | 6.33(1.98)<sup>cde</sup> | 5.70<sup>abc</sup> |
| **Tekisic** | 33.67(3.53)<sup>ab</sup> | 8.50(2.17)<sup>cde</sup> | 6.37<sup>abc</sup> |
| **Yellow Caturra** | 57.50(3.99)<sup>a</sup> | 9.83(2.28)<sup>cde</sup> | 7.47<sup>ab</sup> |
| **SEM (±)** | 0.35 | 0.20 | 0.05 |
| **CV (%)** | 15.33 | 16.63 | 20.17 |
| **LSD** | 0.92 | 0.66 | 1.94 |
| **F-test** | * | *** | * |
| **Grand Mean** | 45.13 | 13.72 | 5.81 |

Note: All mean values with same letters are not significantly different; Duncan's multiple range test; $p < 0.05$ , \*\*\*: significance at 0.1%; \*\*: significance at 1%; \*: significance at 5%; . : significance at 10%; ns: non- significant; dataset inside parenthesis () represents transformed data, CV: Coefficient of Variation; LSD: Least Significant Difference; SEM: Standard error of mean; CV% and LSD are of transformed data.

#### Fruit Morphology

The cherry length across the different genotypes ranges from 12.74 mm to 17.49 mm, with a grand mean of 14.81 mm. Pacas produce the longest cherry (17.49 mm), followed by pacamara (16.59), while Arghakhachi Local recorded the shortest cherry length (12.74 mm). The genotypes Catimor (15.7 mm) and Selection-10 (15.77 mm) are statistically at par in measurement.

The variety Pacas (17.49) recorded the highest cherry length, followed by Pacamara (16.59), whereas Arghakhachi Local recorded the lowest cherry length. The cherry length value ranges from 12.74 to 17.49. The variety Catimor (15.7) and Selection-10 (15.77) were statistically at par.

Regarding the cherry diameter, the variety Indonesia (13.38) produced the largest diameter, followed by Selection-10 (12.85) and Pacamara (12.41). The genotypes Chhetradeep (12.28) and Pacas (12.28) also produced large diameters and were significantly similar. Among 19 genotypes, cherry diameter ranged from 13.38 to the lowest of Bourbon Amarillo (9.40).

**Table 3.**
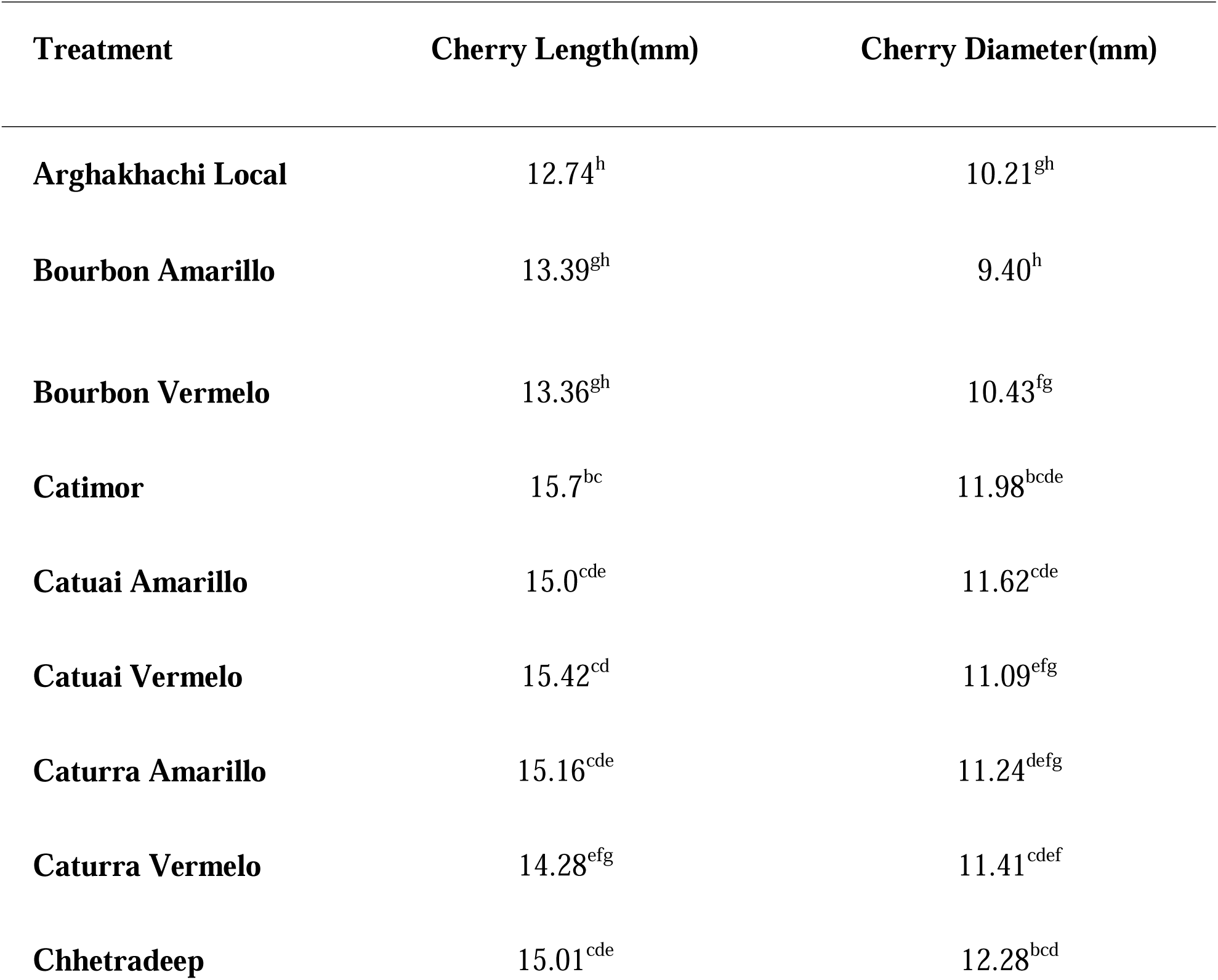
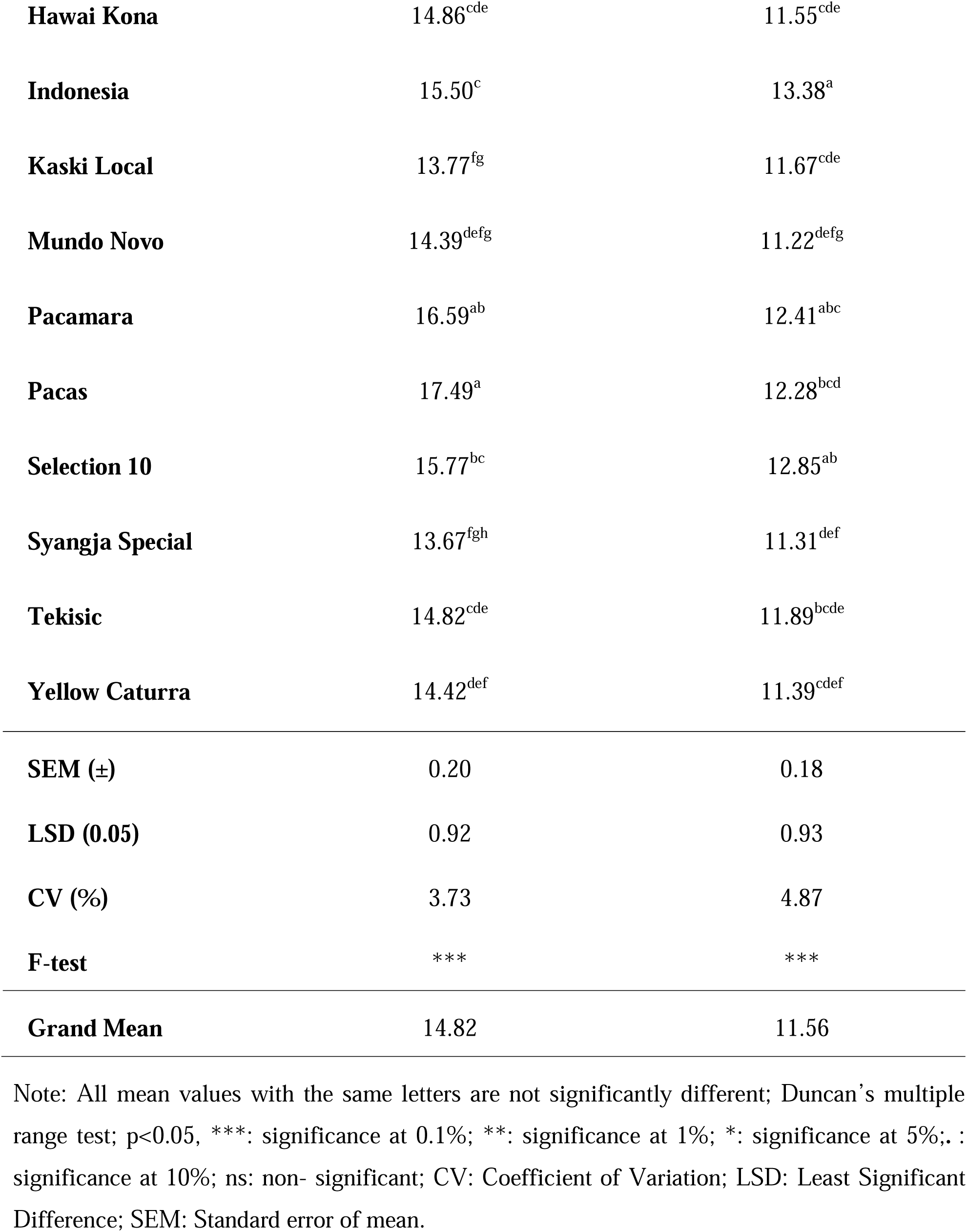
Cherry size of coffee (*Coffea arabica* L.) as influenced by genotypes in Kaski, Nepal, 2026.

#### Yield Parameters

##### Cherry Numbers

Productivity in terms of cherry numbers showed that Catimor (1,840.50) and Yellow Caturra (1,778.80) were the top performers, both exceeding the grand mean of 1,305.82. The lowest cherry count was recorded in Pacas (247.00).

##### Hundred Cherry Weight

The hundred weight of cherry showed a grand mean of 137.81 gm. Both Pacamara and Selection-10 recorded the maximum weight of 167.67 gm. The lowest weight observed in Bourbon Amarillo was 120.33 gm.

##### First-harvest fresh cherry yield (kg plant□¹)

Yellow Caturra recorded the highest first-harvest fresh cherry yield (2.75 kg plant□¹), whereas Catuai Amarillo recorded the lowest, at 0.53 kg plant□¹, statistically at par with Bourbon Vermelo , at 0.54 kg plant□¹.

##### Second harvest yield (kg plant□¹)

The variety Yellow Caturra again showcased the highest yield (2.59 kg plant□¹), which was statistically similar to Catuai Amarillo (2.57 kg plant□¹), followed by Catuai Vermelo (2.20 kg plant□¹). The variety Bourbon Vermelo (0.41 kg plant□¹) was found to be the least yielding.

##### Total yield (kg plant□¹)

Yellow Caturra recorded the highest total yield (5.34 kg plant□¹), followed by Caturra Amarillo (3.56 kg plant□¹). In contrast, Bourbon Vermelo recorded the lowest total yield (0.95 kg plant□¹) among the genotypes.

**Table 4.**
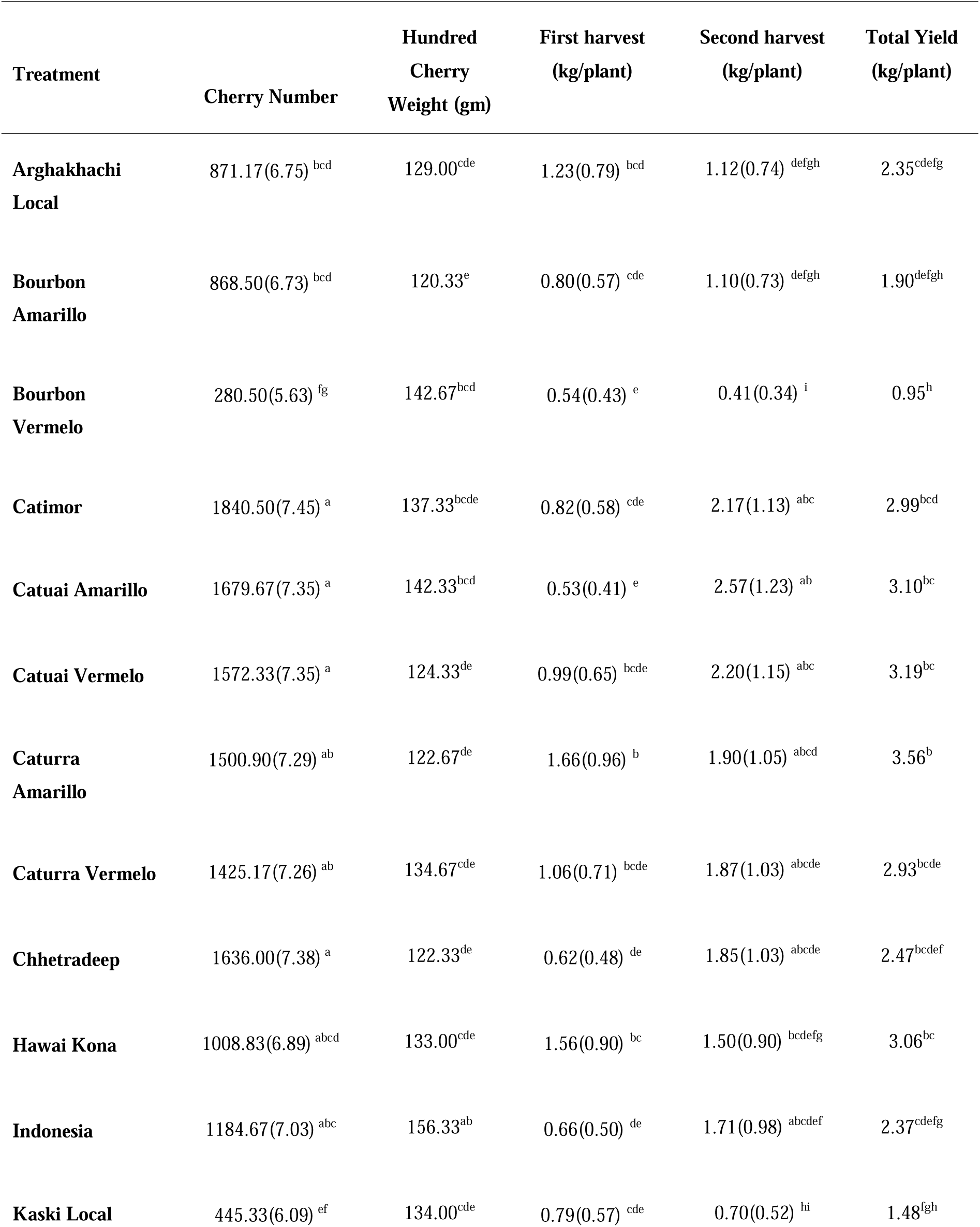
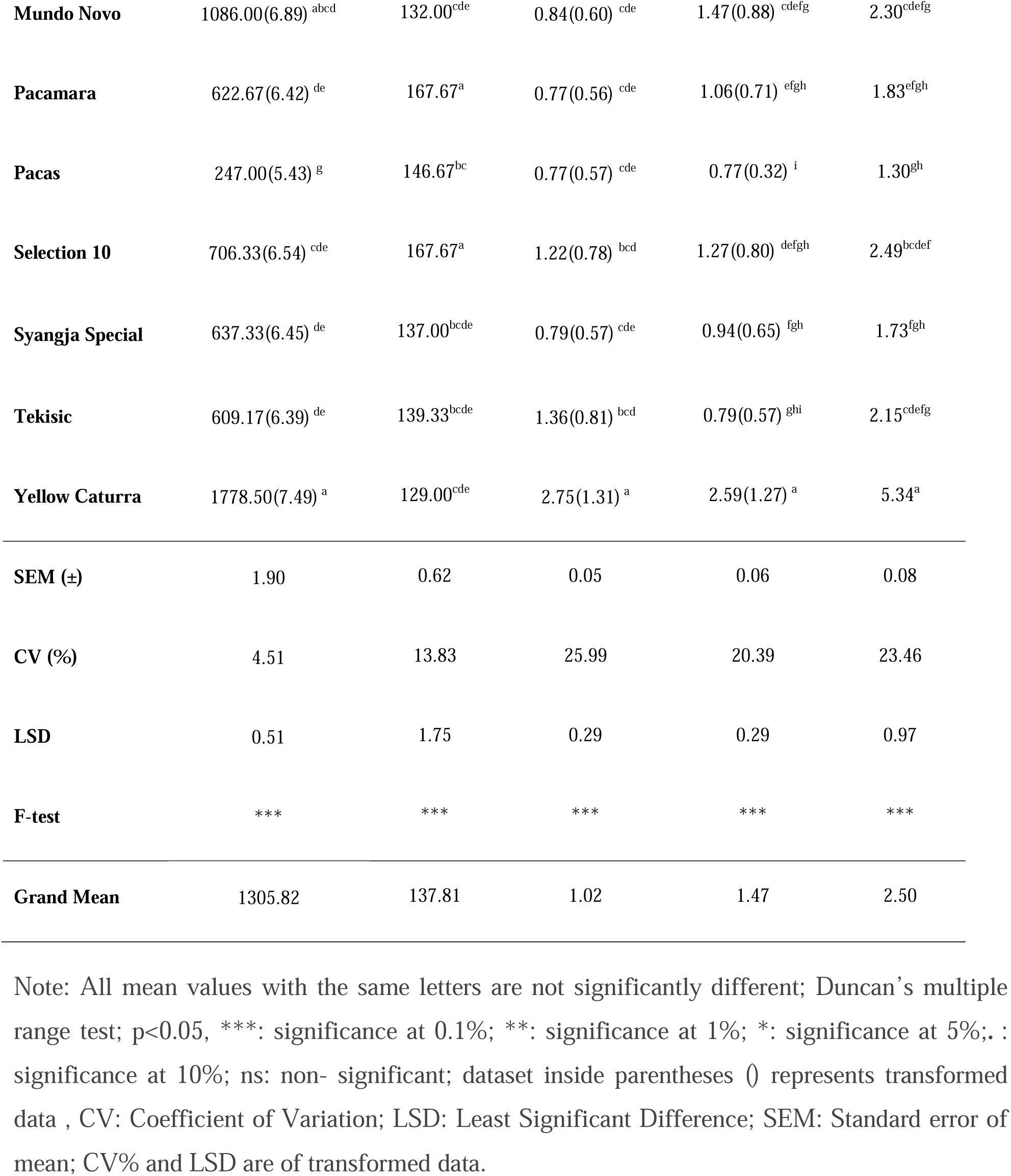
Yield parameters of coffee (*Coffea arabica* L.) as influenced by genotypes in Kaski, Nepal, 2026.

##### Quality Characteristics

The TSS of pulp measured in degree Brix ranged from 25.97 to 16.80 across the genotypes. The variety Catuai Amarillo exhibited the highest TSS concentration, while Tekisic showed the lowest among all. Other genotypes, performing above the mean, included Caturra Vermelo (23.47) and Kaski Local (23.17).

**Table 5.** Quality parameter of coffee (*Coffea arabica* L.) as influenced by genotypes in Kaski, Nepal, 2026.

| Treatment | TSS of pulp (°Brix) |
| --- | --- |
| Arghakhachi Local | 22.30<sup>abc</sup> |
| Bourbon Amarillo | 22.70<sup>abc</sup> |
| Bourbon Vermelo | 19.53<sup>bcde</sup> |
| Catimor | 18.67<sup>cde</sup> |
| Catuai Amarillo | 25.97<sup>a</sup> |
| Catuai Vermelo | 22.43<sup>abc</sup> |
| Caturra Amarillo | 22.10<sup>abcd</sup> |
| Caturra Vermelo | 23.47<sup>ab</sup> |
| Chhetradeep | 19.70<sup>bcde</sup> |
| Hawai Kona | 20.57<sup>bcde</sup> |
| Indonesia | 17.80<sup>de</sup> |
| Kaski Local | 23.17<sup>ab</sup> |
| **Mundo Novo** | 23.00<sup>abc</sup> |
| **Pacamara** | 21.90<sup>abcd</sup> |
| **Pacas** | 18.67<sup>cde</sup> |
| **Selection 10** | 21.57<sup>bcd</sup> |
| **Syangja Special** | 21.17<sup>bcd</sup> |
| **Tekisic** | 16.80<sup>e</sup> |
| **Yellow Caturra** | 22.10<sup>abcd</sup> |
| **SEM (±)** | 0.24 |
| **CV (%)** | 10.48 |
| **LSD** | 3.69 |
| **F-test** | ** |
| **Grand Mean** | 21.24 |

Note: All mean values with the same letters are not significantly different; Duncan's multiple range test; $p < 0.05$ , \*\*\*: significance at 0.1%; \*\*: significance at 1%; \*: significance at 5%; .: significance at 10%; ns: non-significant; CV: Coefficient of Variation; LSD: Least Significant Difference; SEM: Standard error of mean.

**Table 6.**
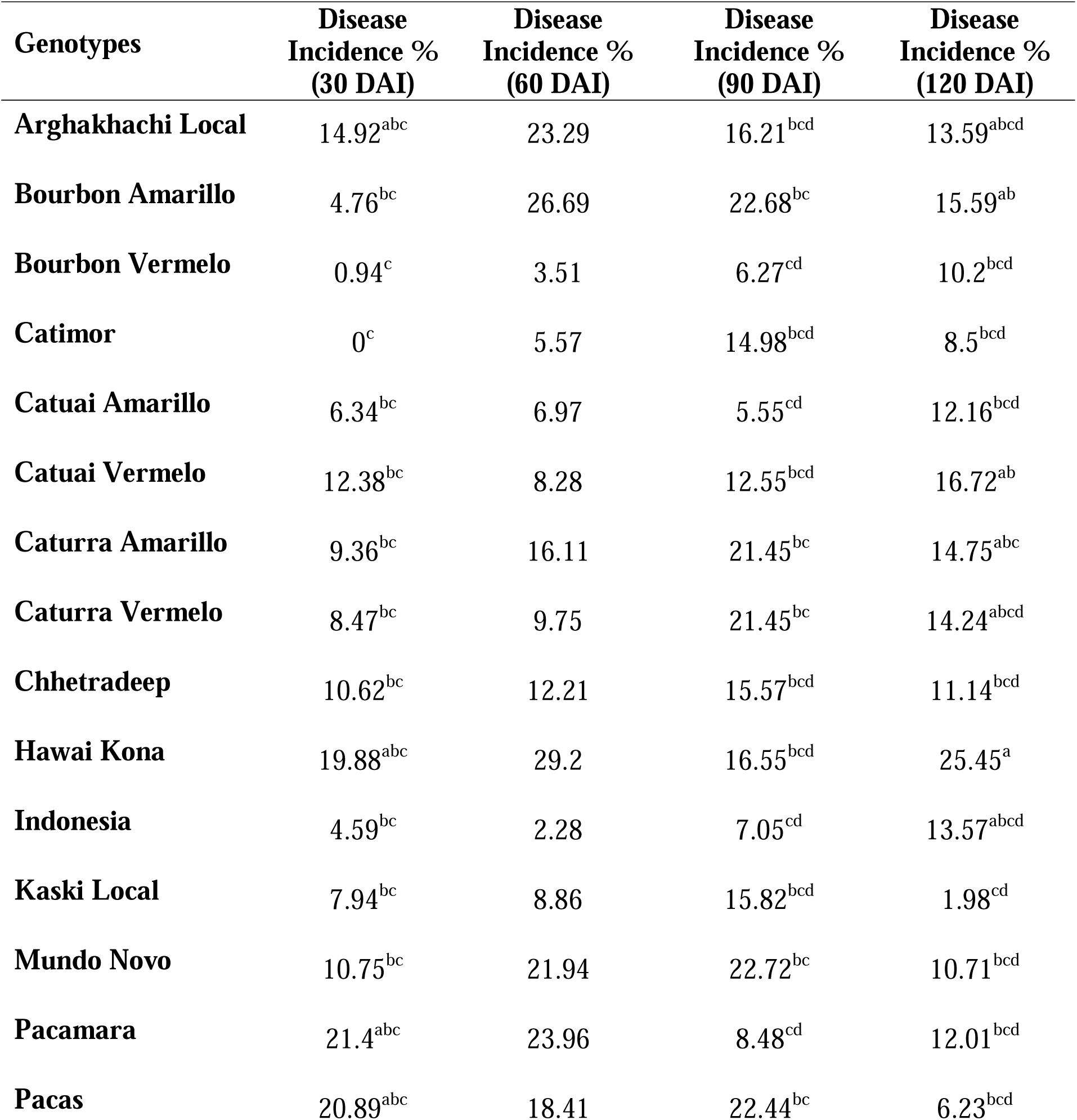
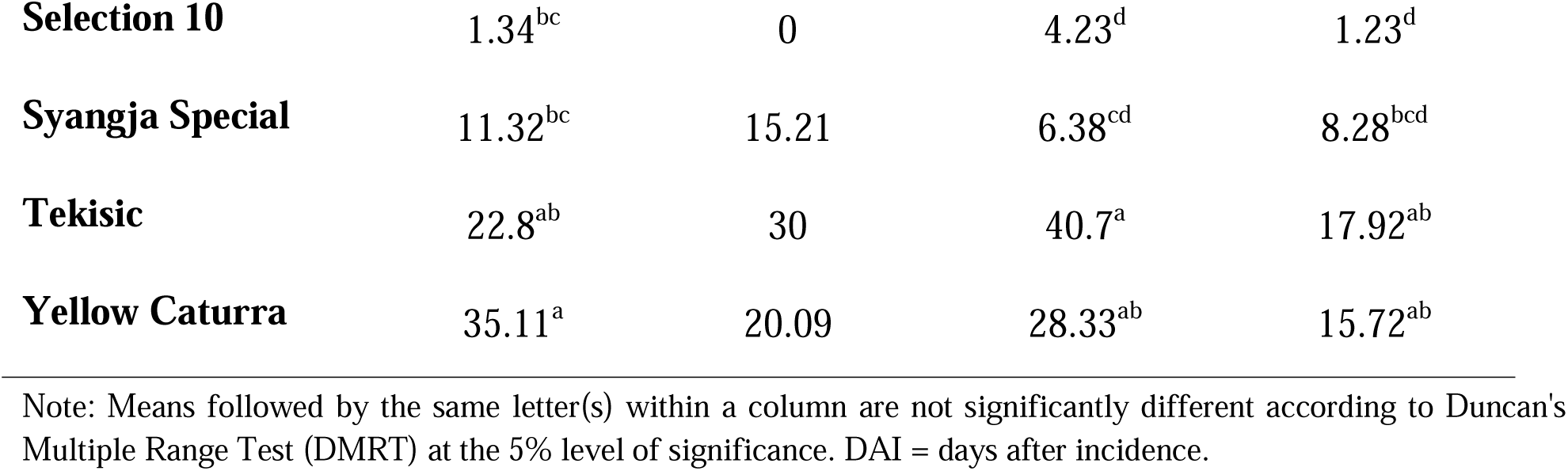
CLR incidence in coffee (*Coffea arabica* L.) as influenced by genotypes in Kaski, Nepal, 2026.

| Genotypes | Disease Incidence %<br>(30 DAI) | Disease Incidence %<br>(60 DAI) | Disease Incidence %<br>(90 DAI) | Disease Incidence %<br>(120 DAI) |
| --- | --- | --- | --- | --- |
| **Arghakhachi Local** | 14.92<sup>abc</sup> | 23.29 | 16.21<sup>bcd</sup> | 13.59<sup>abcd</sup> |
| **Bourbon Amarillo** | 4.76<sup>bc</sup> | 26.69 | 22.68<sup>bc</sup> | 15.59<sup>ab</sup> |
| **Bourbon Vermelo** | 0.94<sup>c</sup> | 3.51 | 6.27<sup>cd</sup> | 10.2<sup>bcd</sup> |
| **Catimor** | 0<sup>c</sup> | 5.57 | 14.98<sup>bcd</sup> | 8.5<sup>bcd</sup> |
| **Catuai Amarillo** | 6.34<sup>bc</sup> | 6.97 | 5.55<sup>cd</sup> | 12.16<sup>bcd</sup> |
| **Catuai Vermelo** | 12.38<sup>bc</sup> | 8.28 | 12.55<sup>bcd</sup> | 16.72<sup>ab</sup> |
| **Caturra Amarillo** | 9.36<sup>bc</sup> | 16.11 | 21.45<sup>bc</sup> | 14.75<sup>abc</sup> |
| **Caturra Vermelo** | 8.47<sup>bc</sup> | 9.75 | 21.45<sup>bc</sup> | 14.24<sup>abcd</sup> |
| **Chhetradeep** | 10.62<sup>bc</sup> | 12.21 | 15.57<sup>bcd</sup> | 11.14<sup>bcd</sup> |
| **Hawai Kona** | 19.88<sup>abc</sup> | 29.2 | 16.55<sup>bcd</sup> | 25.45<sup>a</sup> |
| **Indonesia** | 4.59<sup>bc</sup> | 2.28 | 7.05<sup>cd</sup> | 13.57<sup>abcd</sup> |
| **Kaski Local** | 7.94<sup>bc</sup> | 8.86 | 15.82<sup>bcd</sup> | 1.98<sup>cd</sup> |
| **Mundo Novo** | 10.75<sup>bc</sup> | 21.94 | 22.72<sup>bc</sup> | 10.71<sup>bcd</sup> |
| **Pacamara** | 21.4<sup>abc</sup> | 23.96 | 8.48<sup>cd</sup> | 12.01<sup>bcd</sup> |
| **Pacas** | 20.89<sup>abc</sup> | 18.41 | 22.44<sup>bc</sup> | 6.23<sup>bcd</sup> |
| **Selection 10** | 1.34<sup>bc</sup> | 0 | 4.23<sup>d</sup> | 1.23<sup>d</sup> |
| **Syangja Special** | 11.32<sup>bc</sup> | 15.21 | 6.38<sup>cd</sup> | 8.28<sup>bcd</sup> |
| **Tekisic** | 22.8<sup>ab</sup> | 30 | 40.7<sup>a</sup> | 17.92<sup>ab</sup> |
| **Yellow Caturra** | 35.11<sup>a</sup> | 20.09 | 28.33<sup>ab</sup> | 15.72<sup>ab</sup> |

Note: Means followed by the same letter(s) within a column are not significantly different according to Duncan's Multiple Range Test (DMRT) at the 5% level of significance. DAI = days after incidence.

##### Coffee Dry and Wet Parchment data

Dry parchment weight ranged from 200g to 150g where Hawai Kona recorded the highest (200g) followed by Syangja Special (190g). The lowest dry parchment was 150g recorded on five genotypes, i.e. Arghakhanchi Local, Catimor, Caturra Amarillo, Catuai Amarillo, and Catuai Vermelo.

Genotypes Hawai Kona and Syangja Special recorded the highest wet parchment weight (330g), whereas Catuai Amarillo (280g) recorded the lowest.

##### Coffee Leaf Rust Incidence

CLR incidence varied among genotypes and assessment dates (Table 6). Incidence generally increased or fluctuated during the observation period, with substantial genotype-specific differences at each assessment. Catimor had no incidence at 30 DAI, while Selection-10 recorded no incidence at 60 DAI and remained low at 90 and 120 DAI (4.23% and 1.23%, respectively). Tekisic reached the highest incidence at 90 DAI (40.70%), whereas Hawai Kona recorded the highest incidence at 120 DAI (25.45%). The variation across dates indicates that CLR infection and progression differed considerably among genotypes under the conditions of the trial.

##### Coffee Leaf Rust Severity

Coffee leaf rust (CLR) was recorded in 19 genotypes at 30, 60, 90, and 120 days after incidence (DAI). Mean severity across genotypes rose sharply from 7.49% at 30 DAI to 34.71% at 60 DAI, then increased moderately to 37.58% at 90 DAS, and finally declined slightly to 35.62% by 120 DAI (Figure 1). Selection-10 remained the least affected genotype at every assessment stage, while Arghakhachi Local recorded the single highest severity value observed in the trial, 70.4% at 120 DAI, despite comparatively low severity earlier in the season.

**Figure 1.**
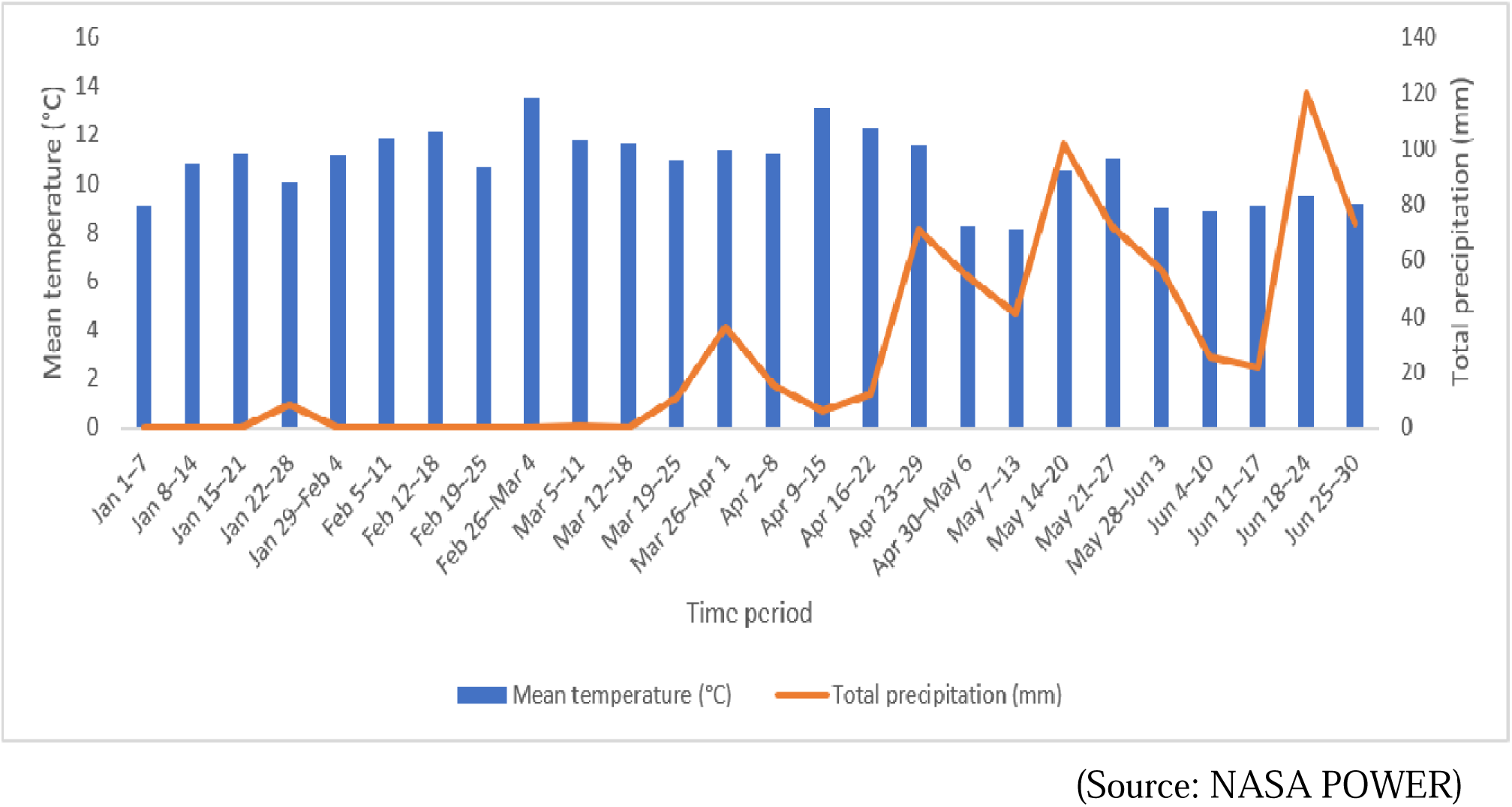
Weekly Temperature and Precipitation during the Crop Growing Period The experiment was conducted in a well-established orchard from which 19 genotypes were selected. A randomized complete block design was used, where genotypes were replicated three times. The research was conducted from January to June 2026.

##### Coffee Rust AUDPC

The area under the disease progress curve (AUDPC) was calculated for each genotype from the four severity readings using the trapezoidal formula of **[25]**. AUDPC ranged nearly 14-fold, from 25.03 rating days in Selection-10 to 347.95 rating days in Caturra Amarillo, around a grand mean of 253.36 rating days (Figure 2). Selection-10 had the lowest AUDPC value, while Caturra Amarillo, Bourbon Amarillo, and Yellow Caturra recorded the highest values.

**Fig 2.**
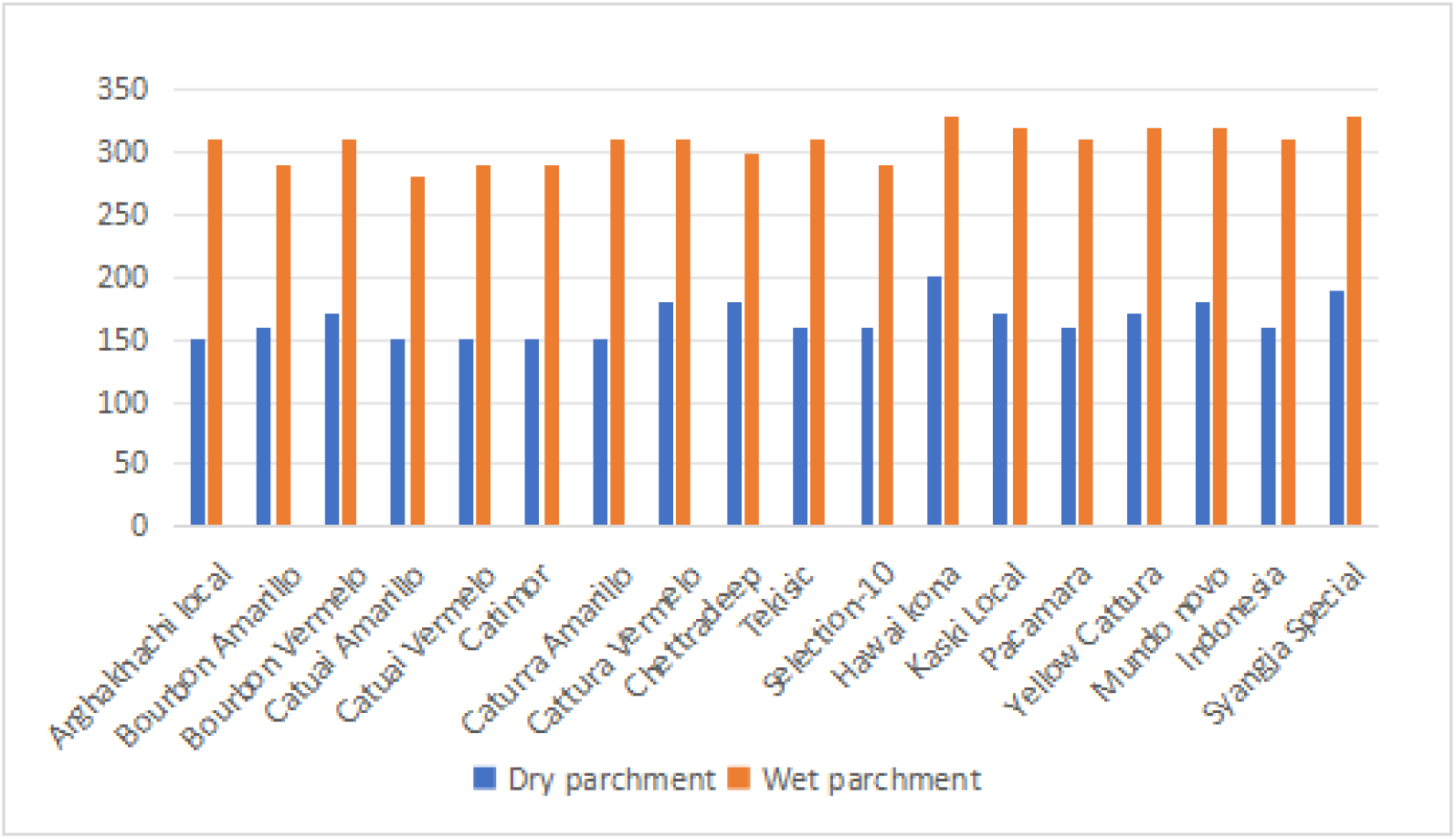
Dry and wet parchment of coffee (*Coffea arabica*) as influenced by genotypes in Kaski, Nepal, 2026

**Figure 3.**
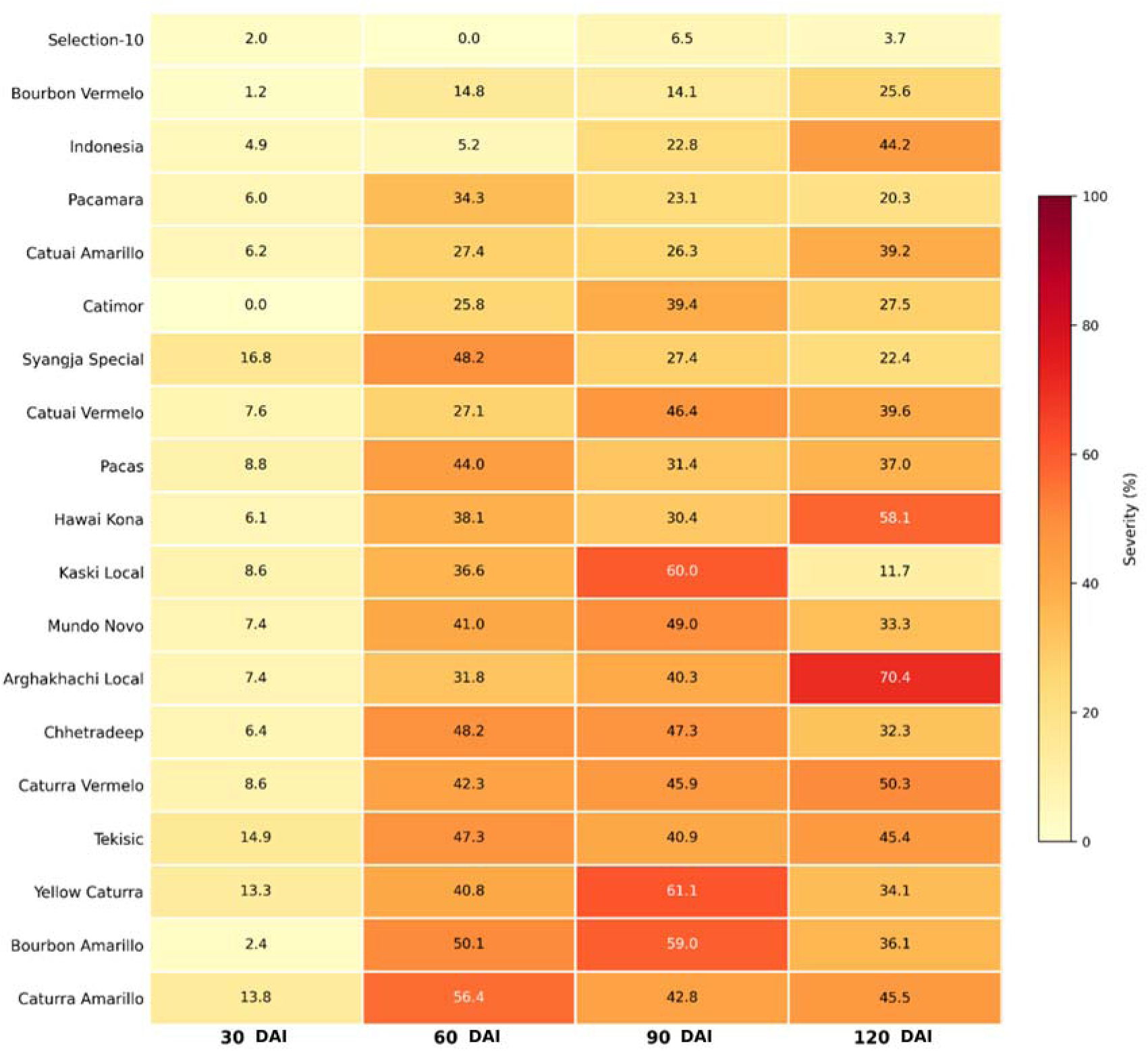
Heatmap of coffee leaf rust severity across 19 coffee genotypes at 30, 60, 90, and 120 DAI in Kaski, Nepal, 2026

**Figure 4.**
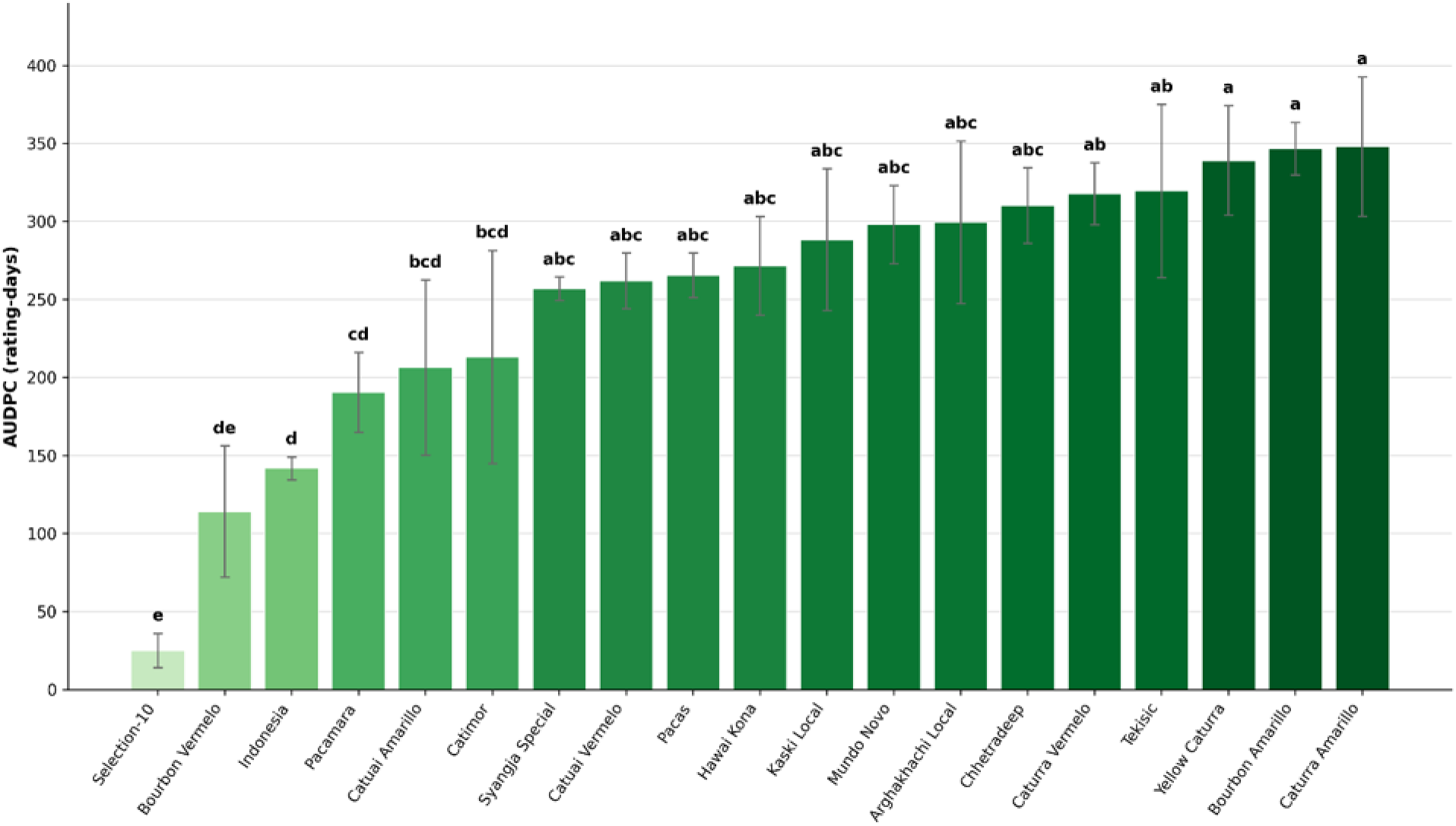
Area under disease progress curve (AUDPC) of coffee leaf rust across 19 coffee genotypes in Kaski, Nepal, 2026

**Figure 5.**
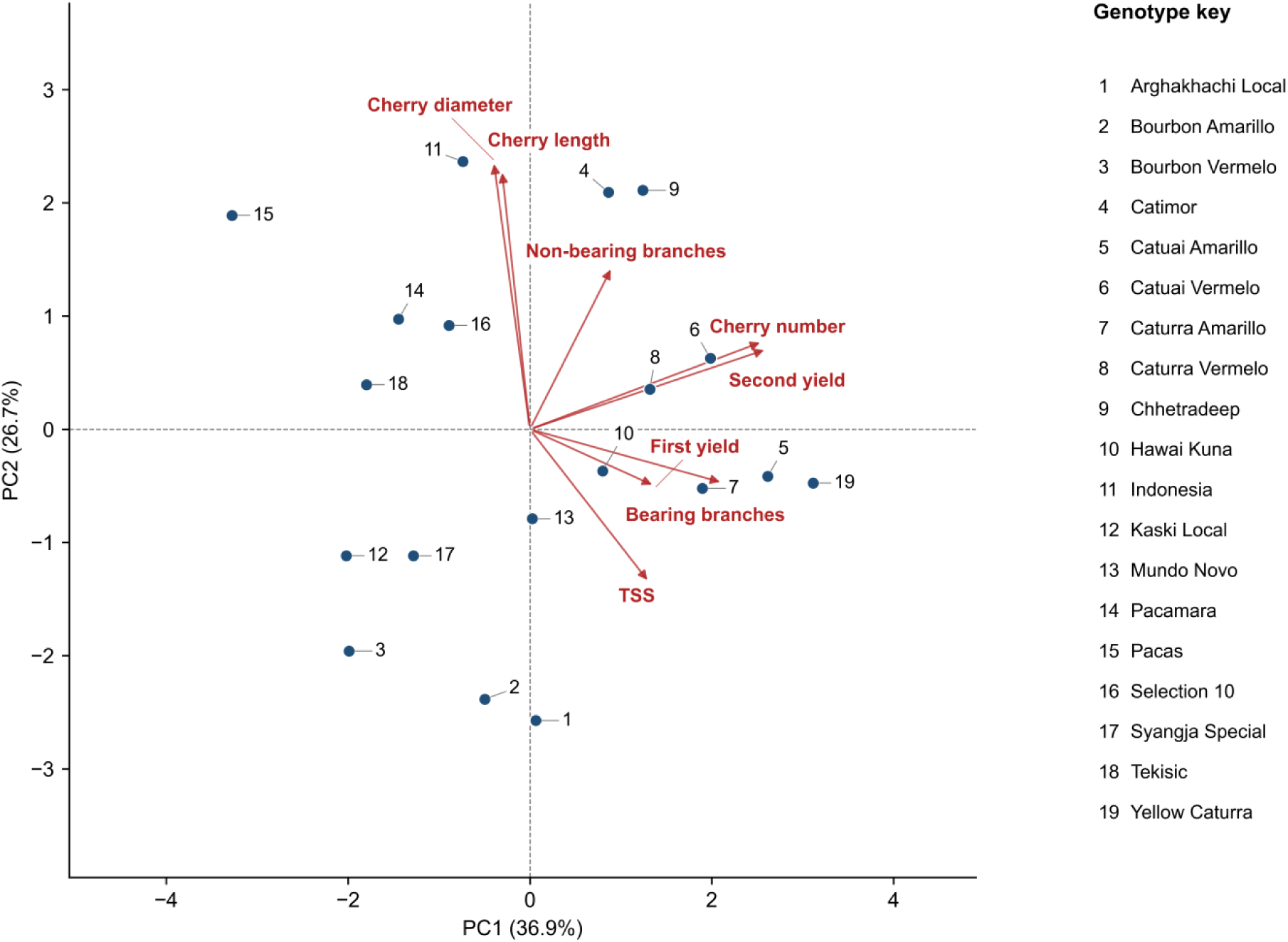
Principal component analysis biplot showing the relationships among coffee genotypes and morphological, yield-related, and quality traits in Kaski, Nepal, 2026.

**Figure 6.**
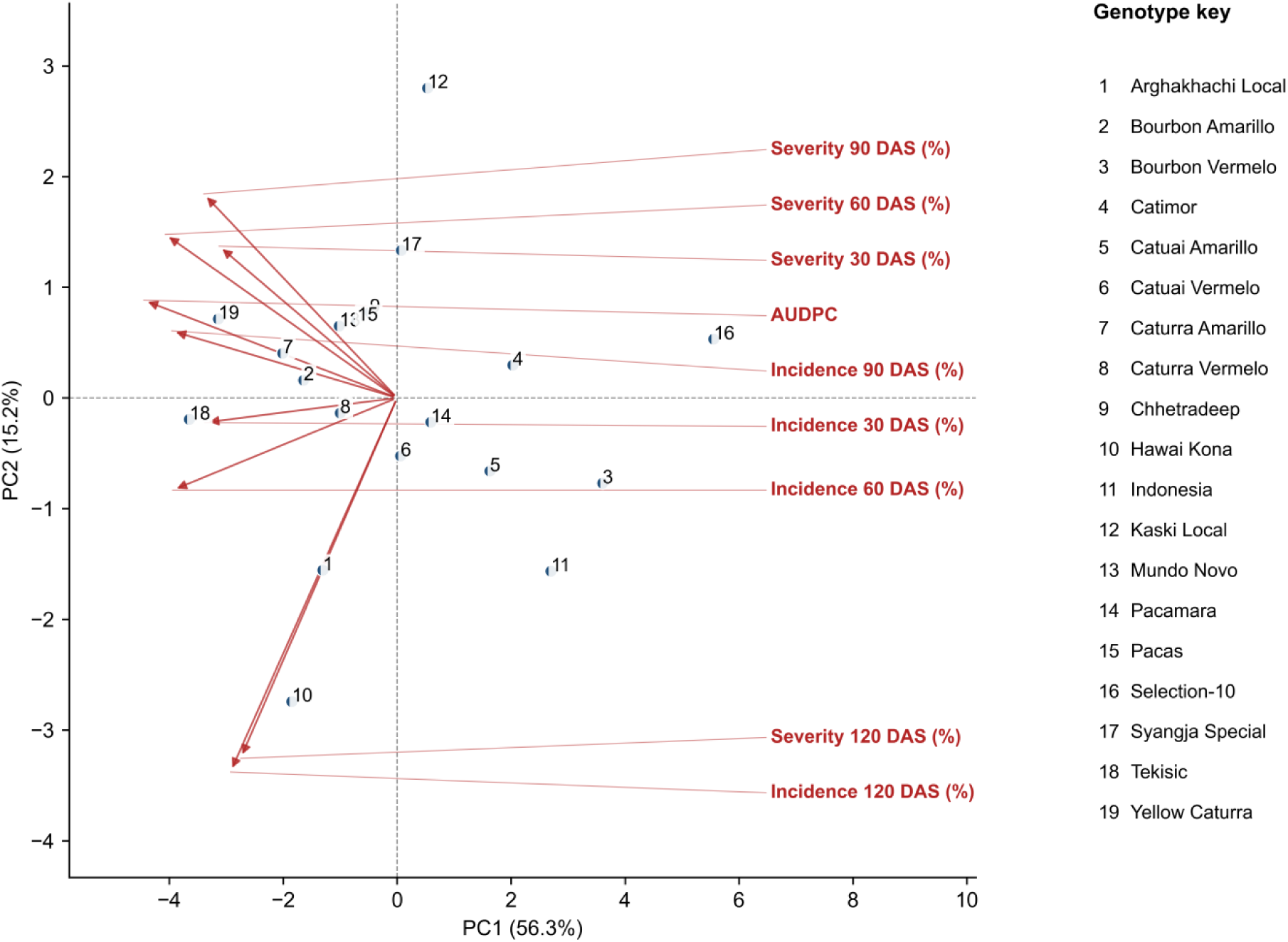
Principal component analysis biplot showing the relationships among coffee genotypes and coffee leaf rust incidence, severity, and AUDPC in Kaski, Nepal, 2026.

The first two principal components explained 63.6% of the total variation among coffee genotypes, with PC1 and PC2 accounting for 36.9% and 26.7%, respectively. Genotypes positioned near particular trait vectors showed corresponding associations with those traits. The separation of genotypes across the biplot indicates substantial variation in agronomic performance and cherry characteristics among the evaluated coffee genotypes.

The first two principal components explained 71.5% of the total variation in coffee leaf rust (CLR) responses, with PC1 accounting for 56.3% and PC2 for 15.2%. Genotypes located in the direction of disease-trait vectors tended to be associated with those responses. Selection-10 was positioned away from several disease-associated vectors, consistent with its comparatively low disease severity and AUDPC in the individual disease assessments. Overall, the biplot illustrates variation among genotypes in the development and progression of CLR under the conditions of the study.

## Discussion

### Morphological traits

#### Number of bearing branches

The number of bearing branches in a coffee plant is a key determinant for coffee yield. Among the 19 genotypes, more than fivefold differences were observed, indicating genotypic control over branching habits, consistent with the report by **[26]**. Caturra Amarillo and Catuai Amarillo, which recorded the highest number of bearing branches in trials, are dwarf-habit genotypes. This compact architecture is also associated with closely spaced secondary branching, a specific trait that is selected to increase fruit-bearing surface within a smaller canopy **[27,28]**. The high bearing branch count observed here is therefore consistent with known branching habits of these genotypes. Hawai Kona also produced a comparably high branch count, but due to the lack of well-established available literature, this result is merely an empirical finding.

#### Number of non-bearing branches

Non-bearing branches are those branches whose fruiting hasn’t started yet; hence, the number of these branches tells about the plant’s vegetative growth. Catuai Amarillo and Caturra Amarillo, recorded the more number of bearing branch, also recorded the lowest non-bearing branch count, which is consistent with the finding of **[29]**, who describe increased fruit number significantly reducing branch number. However, this pattern was not consistent for all the genotypes: Chhetradeep and Caturra Vermelo showed a high number of non-bearing branches despite having a high count of bearing branches, indicating that these genotypes have high overall branching vigor rather than a resource trade-off between bearing and non-bearing structures. This description does not contradict the trade-off principle but indicates the occurrence of a resource trade-off within all the genotypes, but with varying magnitude that depends upon the vigor budget, which varies between genotypes. This interpretation justifies both Chhetradeep and Caturra Vermelo producing a surplus amount of bearing and non-bearing branching. This interpretation reconciles **[29]** with **[30]**, who reported lateral branch number as a genetically variable trait that can correlate positively with yield-related traits across genotypes.

#### Number of cherries per node

The number of cherries per node is one of the key determinants of Arabica coffee yield and has been reported to positively affect clean coffee yield **[31]**. Among the tested genotypes, Caturra Amarillo recorded the highest number of cherries per node (7.73), which is statistically at par with Yellow Caturra (7.47) but superior to most of the other indicated genotypes. The two Catuai genotypes, Catuai Amarillo (6.53) and Catuai Vermelo (6.47), Selection-10 (6.40), Tekisic (6.37), and Caturra Vermelo (6.33) also performed well. In contrast, Kaski Local (4.20), Pacas (4.37), Mundo Novo (4.53), and Bourbon Amarillo (4.93) produced the fewest cherries per node. This superiority of Caturra and Catuai types is consistent with their genetic background, as both of them are compact, short-internode mutants of Bourbon known for dense branching and high fruit-bearing capacity per unit length of branch, therefore selected specifically for high productivity **[27,28]**.

#### 4.2.2 Yield Parameters Fresh cherry yield

Yellow Caturra’s consistent top-ranking yield suggests a stable, high-yielding phenotype insensitive to seasonal fluctuations, consistent with Caturra’s established reputation as a vigorous producer, historically selected as a dwarf-bourbon mutant for short internodes and high node density **[32]**. The yellow cherry phenotype does not appear to compromise this yield potential **[33]**. Catuai Amarillo’s comparable performance also fits its breeding history: the IAS program combined Caturra’s compact habit with Mundo Novo’s bigger size specifically to produce high-yielding, high-density compatible cultivars **[34]**. **[35]** also recorded Yellow Caturra as the highest yielder (3.34 kg/plant) in a Gulmi trial, further supporting its yield edge across sites and seasons in Nepal.

Bourbon Vermelo and Catimor’s poor performance also has a plausible genetic explanation. Bourbon is inherently tall-growing and low-yielding at standard density. A Bourbon Vermelo likely represents a less improved landrace, while Catimor’s low yield despite its rust resistance is consistent with the well-documented yield penalty associated with the rust resistance associated with rust breeding, a resistance that is increasingly becoming prone to breakdown as new races of rust emerge **[36]**.

##### Number of cherries per plant

Catimor (1,840.5 cherries) and Yellow Caturra (1,778.8 cherries) both exceed the grand mean cherry count of (1,305.8 cherries), indicating high fruit-setting capacity in both. The Catimor inconsistency between the high cherry numbers and low overall yield points to individual cherry weight or marketable fruit proportion as a limiting factor and also suggests fruit count alone as a proxy for poor productivity **[37]**. Excessive cherry load without proportional yield gain is consistent with source-sink imbalance affecting dry-mass partitioning **[38]**(Leon Burgos et al., 2024). The same pattern appears in Nepal; **[39]** found that cherry recovery in Catimor and Yellow Caturra depends more on fruit quality than cherry number, and **[35]** found genotypes differ significantly in dry/fallen cherry count, meaning much of a high cherry load can be lost before becoming usable yield.

##### Hundred Cherry Weight

Pacamara and Selection-10 both reached the maximum hundredweight, that is, 167.67 g, which is higher than the mean hundredweight (137.81 g). Pacamara’s large bean size is inherited from its Maragogipe parent **[40]**, which is commercially relevant as bean size is recognized as one of the market quality attributes **[41]**. Selection 10’s large cherry may reflect its Ethiopian Tafarikela parentage, previously linked to above-average crop quality components **[42]**.

Bourbon Amarillo, which has the lowest hundred cherry weight, is consistent with its poor yield performance and cherry number count. The inverse relationship between a high cherry count and lower individual cherry weight, as in Catimor, reflects documented source-sink limitation, indicating a high fruit load reduces assimilate mobilization to individual fruit **[29]**.

##### Size Parameters

###### Cherry length

Cherry length and cherry diameter are key parameters crucial for post-harvest processing and market **[43]**. Pacas and Pacamara exhibiting longer cherry length suggest that these genotypes possess inherently larger fruit architecture than the short-fruited genotypes such as Arghakhachi Local, Bourbon Amarillo, and Bourbon Vermelo, consistent with reports that fruit size traits are strongly genotype-dependent and remain significantly differentiated across the evaluated coffee genotypes **[40]**. The distinct physical variation observed between local landraces (like Arghakhachi Local) and improved mutants (like Pacamara) in this trial mirrors the findings of **[9]**, who also reported significant variations in the physical attributes of coffee genotypes across different Nepalese landscapes. The statistically similar performance of Catimor and Selection-10 for cherry length also fits the pattern that some genotypes form overlapping statistical groups despite overall genotypic-level variability.

###### Cherry diameter

Indonesia exhibited the greatest cherry diameter, while Selection-10, Pacamara, Chhetradeep, and Pacas also produce relatively broad cherries. Previous studies report that larger fruit width is positively associated with larger bean dimensions and 100-bean weights **[40]**, who observed a positive association between cherry length and diameter, noting that the strength of this association differed across genotypes. Pacamara and Pacas, the two longest-cherry genotypes in this trial, were also among those with the largest cherry diameters, supporting the claim. This relationship did not hold uniformly across all genotypes, as Indonesia, which recorded the greatest cherry diameter, did not exhibit a correspondingly long cherry. Such an exception indicates that although cherry length and diameter tend to co-vary, the two traits are not entirely governed by the same genetic mechanisms, allowing some genotypes, such as Indonesia, to develop greater diameter independent of length. These genotype-specific inconsistencies highlight the importance of measuring cherry length and diameter as distinct parameters during varietal evaluation, rather than treating one dimension as a reliable indicator of the other highlighting the need to measure both dimensions independently, a necessity supported by the genetic diversity of Nepalese coffee **[11]**.

##### Quality Parameters

TSS across the 19 evaluated genotypes ranged widely, from 16.80 °Brix in Tekisic to 25.97 °Brix in Catuai Amarillo, with a grand mean of 21.24 °Brix and a moderate CV (10.48%), pointing to genotype as the dominant source of variation rather than environment, since all genotypes were grown and sampled under the same field conditions. This pattern agrees with earlier findings that arabica genotypes differ markedly and heritably in sugar-related and cup-quality traits, with genetic background often explaining more of the variance than site or seasonal effects **[44,45]**. Molecular diversity work on Ethiopian commercial cultivars has also confirmed that named arabica genotypes are genetically distinct enough to justify such trait-level divergence **[46]**. Catuai Amarillo’s strong TSS performance mirrors reports elsewhere that Catuai-type genotypes tend to accumulate soluble solids more efficiently than other commercial types **[47]**, while the comparatively low values in Tekisic, Indonesia, and Catimor reflect the same genotype-driven inefficiency documented in other varietal screening trials.

It is worth noting that TSS reflects sugar status mainly in the cherry pulp and mesocarp rather than the bean itself, since sugar dynamics in these tissues are regulated somewhat independently **[48]**. That fruit ripeness stage can shift TSS readings even within a single variety **[49]**. Because harvesting and processing were standardized across treatments here, such maturity-related noise was minimized, reinforcing that the differences observed are largely varietal. Taken together, the above-mean performance of Catuai Amarillo, Caturra Vermelo, and Kaski Local marks them as promising candidates for sweetness- and quality-oriented selection in future coffee improvement.

##### Disease traits

###### Disease incidence

Selection-10 maintained the lowest coffee leaf rust incidence throughout the entire screening window, consistent with evidence that some Ethopian derive germplasm express intermediate to partial resistance and can remain less affected across environments **[50,51]**. This prolonged suppression of infection establishment suggests durable partial or polygenic resistance **[52,53]**. GxE interaction is also important in CLR expression, so a stable low-incidence genotype across the whole screening window is especially valuable as breeding material **[54]**.

Similarly, Catimor displayed a complete absence of CLR at 30 DAI and low early incidence at 60 DAI, but incidence progressively increased at 90 DAI. The initial resistance shown by Catimor may be due to the presence of S_H_6, S_H_7, S_H_8, and S_H_9 resistance genes that are derived from the Timor hybrid (*C. canephora* x *C. arabica*) **[55–57]**. However, the rise in Catimor incidence during later stages is also consistent with well documented erosion of S_H_ based resistance under field selection pressure as new strains of CLR emerges **[51,56,58]**. This limitation is not unique to Catimor, as many cultivar with vertical resistance lose their effectiveness after years of exposure, whereas broader pyramiding of partial resistance tends to be more durable approach **[59,60]**.

Dwarf pure Arabica genotypes, such as Yellow Caturra and Tekisic, exhibited rapid initial and peak disease initiation incidence, because the non-introgressed pure Arabicas are generally more susceptible than interspecific lines carrying rust resistance genes **[50,59]**.

###### Disease severity

Across all 19 genotypes, mean leaf rust severity increased sharply from 7.49% at 30 DAI to 34.71% at 60 DAI, peaked at 37.58% at 90 DAI, and stabilized at 35.62% by 120 DAI. This pattern of disease severity reflects the sigmoid curve progression that mimics the polycyclic nature of CLR **[61–63]**. The severity pattern also matches reports that CLR severity and incidence often rise seasonally to peak during favourable periods, including harvest, post-monsoon, or wet season windows **[50,64–66]**.

Selection-10 remained the least affected genotype across all screening dates, suggesting it successfully restricted lesion expansion and limited the secondary sporulation **[52,54]**. In contrast, Arghakhachi local recorded the highest severity value despite showing relatively lower severity earlier in the season. This late-season surge demonstrates a lack of durable adult-plant horizontal resistance, because some cultivars initially appear less affected but lose effectiveness under repeated field infection cycles **[51,59]**.

###### AUDPC

Selection-10 had the lowest AUDPC value throughout the screening period, showing season-long field resistance, as AUDPC captures both the delayed epidemic onset and reduced disease progression over time **[54,61,66]**. When combined with its superior hundred weight and high pulp, it emerges as a valuable germplasm.

Conversely, Caturra Amarillo, Bourbon Amarillo, and Yellow Caturra recorded the highest AUDPC values in the study, indicating their extreme epidemiological vulnerability under subtropical Nepalese conditions, which also agrees with the broader susceptibility of traditional Arabica backgrounds **[50,53,64]**. Despite its high AUDPC value, Yellow Caturra produced the highest fresh cherry weight in the trial. This divergence between high disease severity and high yield points towards short-term disease tolerance in Yellow Caturra, which is driven by its high node density and fruit-setting capacity. However, the literature suggests that higher unmanaged yield loads on highly rust-susceptible genotypes increase assimilate demands, drawing phenolic defense compounds away from leaves toward the developing fruits, causing source-sink imbalance, leaving plant foliage defense compromised, escalating the risk of defoliation, and resulting in severe yield collapse in the upcoming cropping cycle **[19,53,57,63]**.

## Conclusion

This study assessed 19 *Coffea arabica* genotypes, grown under a Randomized Complete Block Design with three replications at the Horticulture Research Station, Kaski, Nepal, with the aim of evaluating their yield performance and pinpointing genotypes with superior yield, morphological, and quality traits.

The genotypes differed significantly across yield and related traits, leading to rejection of the null hypothesis. Yellow Caturra produced the greatest total fresh cherry yield (5.34 kg/plant), trailed by Caturra Amarillo (3.56 kg/plant), Catuai Vermelo (3.19 kg/plant), and Catuai Amarillo (3.10 kg/plant), whereas Bourbon Vermelo and Catimor gave the poorest yields. This pattern traced back to underlying morphological differences: bearing branch number varied more than fivefold across genotypes, peaking in Catuai Amarillo (76.00) and Caturra Amarillo (72.00), while cherries per node were greatest in Caturra Amarillo (7.73) and Yellow Caturra (7.47) traits that directly explain their yield advantage.Weighing yield alongside morphological and quality traits singled out Yellow Caturra and Caturra Amarillo as the strongest overall performers, both compact, densely-noded types well matched to Nepal’s terrace-farming systems. On the quality side, Pacas and Pacamara recorded the longest cherries, and Pacamara together with Selection-10 produced the heaviest hundred-cherry weight (167.67 g), pointing to specialty-market suitability; Catuai Amarillo further stood out with the highest total soluble solids (25.97°Brix), signaling strong cup-quality potential.

Collectively, these results demonstrate that variety choice strongly shapes coffee productivity and quality in Kaski district and can be matched to different production goals Yellow Caturra and Caturra Amarillo for maximizing yield, Catuai Amarillo and Catuai Vermelo where both yield and quality matter, and Pacamara and Selection-10 for premium markets, with Catimor still valuable in areas where rust pressure is high despite its lower yield. Since this trial covered only one season and one location, these results should be viewed as a preliminary screening rather than definitive recommendations. Broader trials spanning multiple seasons and locations, paired with cup-quality testing and rust-resistance evaluation, are needed to confirm these findings and guide reliable varietal recommendations for Nepal’s growing coffee industry.

